# A conserved Rhs module associated with the type VI secretion system facilitates effector innovation and immunity acquisition for ecological adaptation of *Stenotrophomonas* species

**DOI:** 10.64898/2026.07.13.738213

**Authors:** Boris Taillefer, Agathe Brault, Alain Sarniguet

## Abstract

The type VI secretion system (T6SS) is a widespread antibacterial weapon whose evolutionary flexibility promotes bacterial survival in competitive environments. Here, we characterized the *vgrG6* cluster of *Stenotrophomonas rhizophila* CFBP13503, which encodes a putative PAAR-Rhs-fused effector (Rhs-Ct, Tse6) along with a poly-immunity cluster. Comparative genomics across *Stenotrophomonas* genus revealed strong conservation of the *vgrG6* core genes, contrasting with a striking variability in the downstream poly-immunity region, consistent with rapid diversification and niche-specific adaptation. *In silico* structural analysis of Rhs-Ct domains from 158 *Stenotrophomonas* strains identified 14 distinct effector families with diverse putative enzymatic activities, including nucleases and deaminases. The Rhs-Ct from strain CFBP13503, Tse6, harbors a domain of unknown function that is rare across the bacterial diversity. Its closest orthologs were found in Gram-positive *Actinomycetes* and halophilic Gram-negative *Planctomycetia*. Strikingly, saline conditions significantly enhanced both *S. rhizophila* growth and T6SS activity. Functional assays demonstrated that Tse6 is specifically active against the phytopathogen *Clavibacter michiganensis* and the plant beneficial strains *Curtobacterium herbarum* and *Plantibacter flavus*, abolishing their resistance to *S. rhizophila* T6SS attacks. Furthermore, the immunity protein Tsi6 was predicted to interact with Tse6 orthologs from phylogenetically distant taxa, indicating a broad protective capacity. Taken together, our results establish the *vgrG6* cluster as a flexible adaptive module that links effector innovation and immunity diversification to ecological specialization. This work highlights the previously unrecognized role for the *S. rhizophila* T6SS in mediating bacterial competition in saline niches dominated by certain Gram-positive species.

**IMPORTANCE:** *S. rhizophila* CFBP13503 carries a large set of T6SS effectors, some of which are specifically targeting bacterial species. It is important to determine which species are targeted and how effectors adapt to changing environments. *Stenotrophomonas* species possess a conserved *vgrG6* cluster composed of a Rhs-fused effector (Rhs-Ct) and a poly-immunity region. Comparative analysis showed that the Rhs-Ct effector varies from stain together with the poly-immunity content. This diversity underlies the adaptive potential of the vgrG6 cluster in Stenotrophomonas species, notably through targeting of Gram-positive bacteria in a very distinct ecological niche. This study reveals a new effector repertoire to target Gram-positive phytopathogens that can be investigated for biocontrol studies.

**Graphical abstract:** 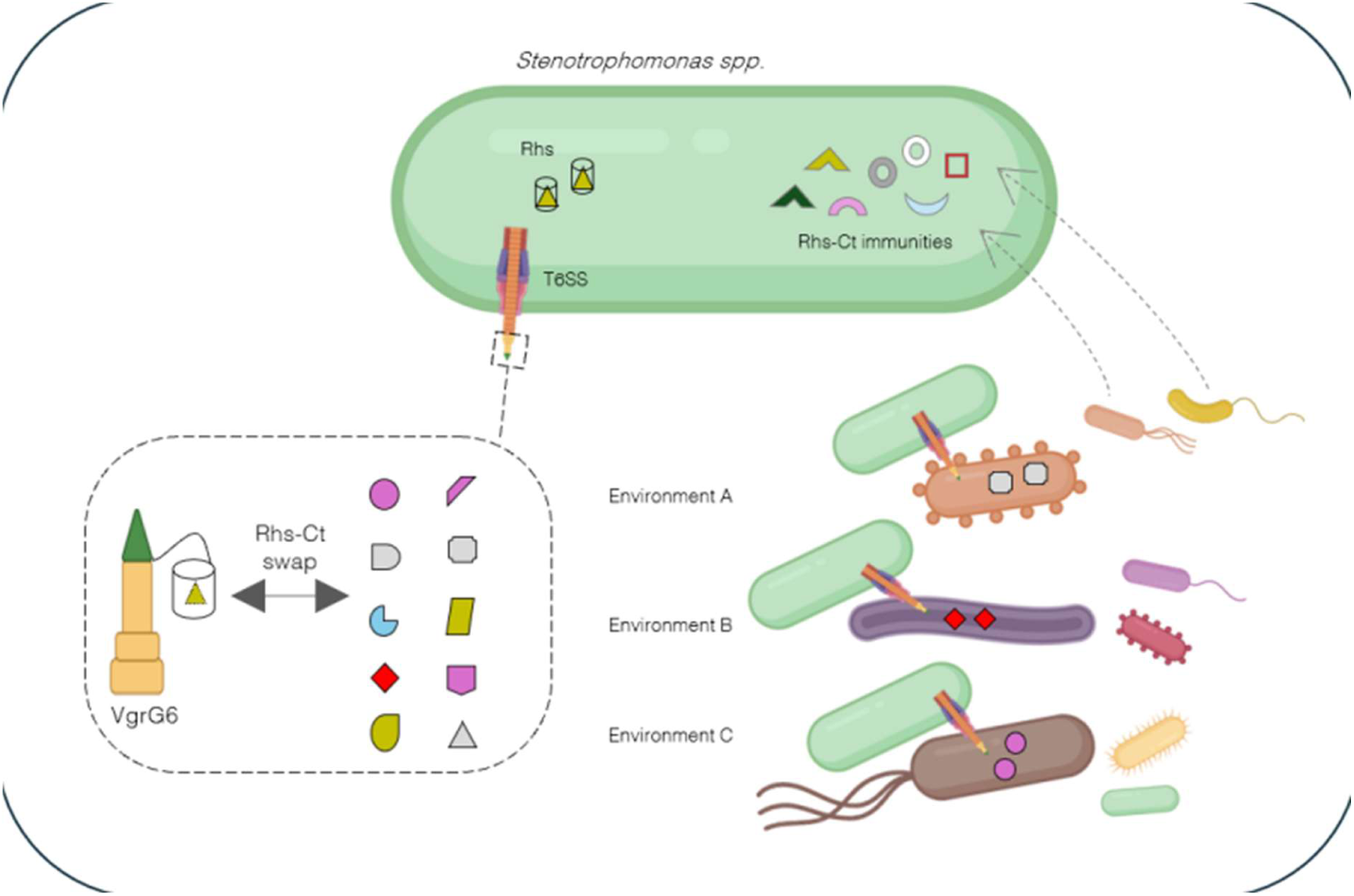

## INTRODUCTION

Bacteria engage in interbacterial competition for space and nutrients to increase fitness in a polymicrobial environment. The type VI secretion system (T6SS) is a contact-dependent weapon that enables bacteria to directly intoxicate neighbouring cells through toxin injection. While T6SS primarily targets both Gram-negative and Gram-positive bacteria, it can also act against eucaryotes as an anti-predatory or virulence mechanism (1, 2). Basically, the T6SS assembles a contractile tail mounted with a puncturing device that is loaded with a toxic effector and propelled upon contraction. This puncturing device is composed of a VgrG trimer spike, which is often reinforced with a metal-bound tip of PAAR. The effector cargo is recruited onto the VgrG through a dedicated type VI adaptor protein (Tap) (3). Effectors are structurally diverse: they can be found as independent or orphan genes, or directly fused with their carrier VgrG, PAAR, Hcp or Rhs as a protein domain. To avoid self-intoxication and stabbing between siblings, the immunity protein (i.e. the cognate anti-toxin) is typically co-expressed with the effector and neutralises its toxic activity (4).

T6SS are genetically encoded in gene clusters spanning 20-80 kbp that carry essential genes for structure assembly and regulation as well as effector-immunity modules. Each module is typically composed of *vgrG*, *paar*, *tap* and effector-immunity pair genes (4). Because the interaction between an effector and its cognate immunity is highly specific, multiple homologous immunity genes are often associated with a single module, providing protection against a range of closely related effectors (5, 6). In addition, non-cognate immunities can be recruited in these clusters to confer wide protection (7–9). Such a poly-immunity cluster, together with their associated effector, are subjected to strong selective pressure from the biotic environment and hence are prone to rapid evolution (10). Consistent with this, numerous comparative genomic studies showed that T6SS genes and associated effectors undergo extensive diversification through gene gain and loss, duplication, mutation and horizontal gene transfer, underlying the central role of effector-immunity pairs in niche adaptation (11, 12).

One striking example of effector diversification involves Rhs-fused effectors encoded within the C-terminal domain of Rhs proteins (Rhs-Ct). The Rhs scaffold -named for rearrangement <u>h</u>ot <u>s</u>pot-is composed of multiple tandem YD-residue repeats domains that facilitate Rhs-Ct exchange through recombination (13–15). Furthermore, these Rhs-Ct are frequently identified as polymorphic toxins, a broadly distributed toxin characterized by extensive diversification in catalytic activity (17–19). This variability within the same species has led to speculation on their potential role in kin discrimination and intraspecific competition (6, 20, 21). However, whether Rhs-Ct variation is involved in speciation and niche adaptation remains poorly investigated. A compelling hint comes from experimental evolution in *Salmonella enterica*, where evolved clones recombined an orphan effector onto an existing Rhs-fused effector, thereby swapping Rhs-Ct catalytic activities and outcompeting ancestral strains in co-culture (22).

The *Stenotrophomonas* genus includes environmental, human-pathogenic and plant-beneficial species, several of which carry at least one T6SS (23). Among them, *S. rhizophila* strains are notable plant-growth promoting bacteria found across diverse environments including plant seeds and tissues, soil, and water (24). *S. rhizophila* CFBP13503 notably deploys its *Sr*-T6SS to compete with seed microbiota and plant pathogens (23, 25). Its high competitiveness in microbial communities stems from a large and diversified effector-immunity repertoire composed of putative DNases, NADases, amidases, phospholipases or pore-forming toxins (26). We recently identified the *vgrG6* module as a candidate weapon against Gram-positive bacteria, based on several distinct genomic features such as a lower GC content relative to the overall *Sr*-T6SS cluster, an unusually long predicted VgrG tip structure, a rare PAAR-Rhs-fused effector of unknown function (Tse6) and an associated poly-immunity gene cluster (26). Intriguingly, the five putative immunity genes share no detectable sequence relatedness with each other, suggesting that each protects against a structurally and functionally distinct effector. Given the rapid evolutionary dynamics of Rhs-fused effectors, whether the *vgrG6* cluster is conserved across *Stenotrophomonas* strains and contributes to niche-specific adaptation remains an open research question.

In this study, we combined comparative genomics, structural prediction, and functional assays to dissect the organization, evolution, and activity of the *vgrG6* cluster in *Stenotrophomonas* spp. Comparative genomics revealed that the core *vgrG6* cluster is highly conserved whereas the toxic Rhs-Ct module and the downstream poly-immunity cluster are highly variable. Using structure prediction and protein-protein interaction modelling (27, 28), we propose that each immunity encoded within the *vgrG6* cluster can specifically inhibit a distinct Rhs-Ct effector found in other bacterial strains. Phylogenetic analysis further showed that Tse6_CFBP13503_ is shared with saline environment-associated *Actinomycetes* and *Planctomycetes*, and that its cognate immunities Tsi6a and Tsi6b are predicted to neutralize Tse6 orthologs. Fluorescence microscopy revealed that saline conditions upregulate *Sr*-T6SS activity while promoting *S. rhizophila* CFBP13503 growth. To assess the anti-Gram-positive potency of Tse6, we performed competition assays against diverse strains isolated from plants. Tse6 was able to inhibit the growth of three Gram-positive targets, among which the phytopathogen *Clavibacter michiganensis*, whereas most of the tested strains were *Sr*-T6SS-resistant. Taken together, our data support the hypothesis of the high adaptive potential of the *Stenotrophomonas*-associated *vgrG6* cluster in competing with Gram-positive bacteria in specific ecological niche.

## RESULTS

### Genetic organization of the vgrG6 cluster indicates a role in competition with Gram-positive bacteria

The 15.4 kb-long *vgrG6* cluster is flanked upstream by the *degQ* operon and downstream by a two-genes operon (**Fig. 1B**). Both operons encode for putative periplasmic peptidase systems S1 or S9 involved in protein quality control or in signalling under specific conditions (29, 30) (<u>Fan2012</u>). The *vgrG6* cluster contains 14 genes in the same genetic orientation, notably *vgrG6* encoding the T6SS spike, *tap6* encoding the effector adaptor, *paar-rhs-tse6* encoding the toxic effector fused with its carrier and two *tsi6a* and *tsi6b* genes encoding the putative immunities. The PAAR-Rhs C-terminal domain (Rhs-Ct) Tse6 shares structural similarities with a *Mycobacterium chelonae* T7SS toxin (DYE20_11250) predicted to act in the cytoplasm as an ADP-ribosyltransferase (**Fig. 1B**). Since we were not able to detect any typical fold or motif linked to ADP-ribosyltransferase activity in Tse6 (31), we will assign it as a nuclease. The cognate T7SS anti-toxin DYE20_11255 is annotated as a pentapeptide repeat-containing protein and possesses structural similarities to Tsi6a (**Fig. 1C**). The downstream region contains diverse immunities genes, thus defining a poly-immunity cluster. Gene *314* encodes a putative LRR domain-containing protein like ribonuclease inhibitors (17). Genes *315*, *317* and *318* all carry a putative Knr4/Smi1 domain characteristic of immunity proteins that neutralize nucleic acid-degrading toxins (14, 20). Gene *315* is the structural orthologue of the YobK antitoxin from the T7SS toxin-antitoxin-II module YobK-YobM of *Bacillus subtilis* or *Bacillus velenenzis* (32, 33) whereas genes *317* and *318* are predicted to interact and together reconstitute a full length YobK-like protein. Intriguingly, *315* but not *317*-*318* displays a signal peptide, suggesting different modes of actions. Gene *316.1* encodes a putative protein with a Imm26 domain predicted to inhibit URI-2 nucleases (17). Gene *316.2* encodes a protein of unknown function with no detectable functional domains, but carrying a signal peptide derived from Gram-positive bacteria. Gene *319* encodes a protein of unknown function that resembles a putative membrane sensor only found in *S. rhizophila* species.

**Fig. 1.**
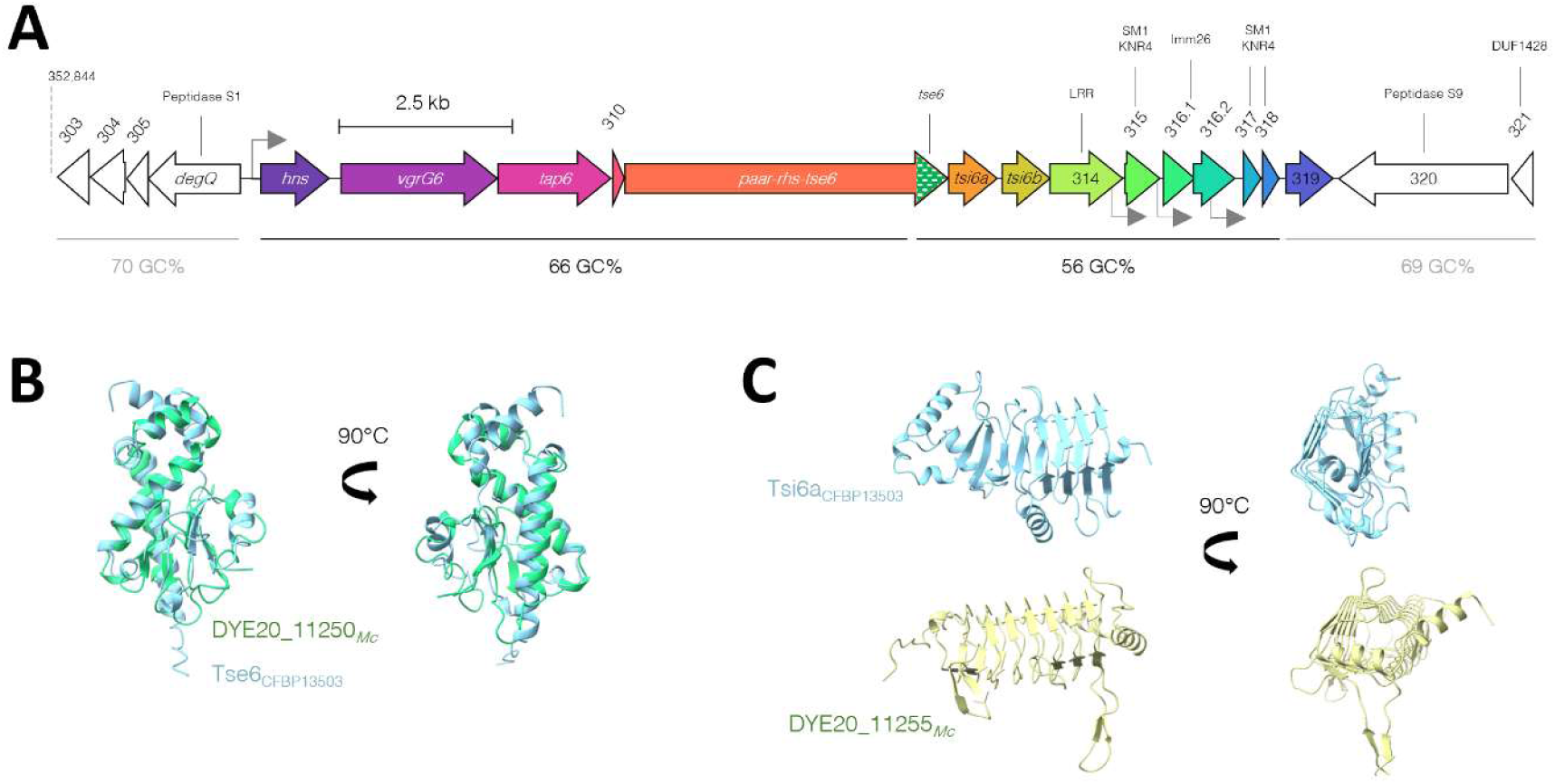
Genetic organization of *vgrG6* cluster and its putative effector Tse6. **A)** Schematic of *vgrG6* genetic organization in *S. rhizophila* CFBP13503. The cluster is composed of 14 genes in the same genetic orientation, from *hns* to 319 genes (colored blocks). The genetic environment of the *vgrG6* cluster upstream and downstream is indicated in white blocks. Gene scale is indicated by the closed line. Putative promoters are indicated by a grey arrow. The GC content of each region is indicated. **B)** Superimposition of Tse6_CFBP13503_ and the putative nuclease DYE20_11250*_Mc_* (*Mycobacterium chelonae*). Structures were predicted by AlphaFold3 and aligned by Matchmaker tools under ChimeraX software. **C)** Comparison of Tsi6a_CFBP13503_ and DYE20_11255, the cognate immunity of DYE20_11250*_Mc_* (*Mycobacterium chelonae*). DYE20_11250*_Mc_* is predicted as a BTB/POZ-containing protein. Structures were predicted by AlphaFold3 and visualized under ChimeraX software.

Based on current mechanistic knowledge of T6SS and Rhs biology, we propose that the chaperone Tap6 contacts the PAAR domain of the fused PAAR-Rhs-Tse6 effector to load it onto the tip of VgrG6 spike that should be secreted directly into a target cell when the T6SS is firing (**Fig. S1**). The Rhs domain encapsulates Tse6, which should be cleaved through two dedicated motifs upon membrane insertion in the target cell. *S. rhizophila* cells produce the cognate Tsi6a immunity to prevent self and sibling-intoxication.

We compared the VgrG6 predicted structure to other anti-Gram-positive VgrGs and found that the RhsB-associated VgrG3 (34) as well as the anti-fungal VgrG5 (35) from *Paracidovorax citrulli* are also predicted to adopt a long tip (**Fig. S2**). The VgrG6 structure and the genetic origin of the poly-immunity cluster reinforce the hypothesis of the role of the *vgrG6* cluster in the competition with Gram-positive bacteria.

### Comparative genomics revealed that *vgrG6* cluster is conserved but the poly-immunity cluster is highly variable in *Stenotrophomonas*

Given the propensity of Rhs effectors to be highly polymorphic and recombine quickly, we hypothesized that both the Rhs-Ct and the downstream poly-immunity region can represent exchangeable adaptive units for niche adaptation (5, 36). To explore this adaptive potential, we performed a comparative genomic analysis of the entire *vgrG6* cluster across the *Stenotrophomonas* genus. Using the Cblaster tool (37), we identified homologous *vgrG6* clusters across 163 genomes out of 193 that possess a T6SS. We observed that *vgrG6*, *tap6*, gene *310* and *paar-rhs-tse* are highly conserved across strains (**Fig. 2A**). Since the VgrG6 tip length may be functionally relevant, we verified that its predicted size is similar in all strains with 23.55 +/-0.37 Å (748.7 +/-1.99 aa). Nonetheless, we found rare adaptations consisting of PAAR domain removal or Rhs domain length reduction. We observed that gene *319* is mostly restricted to *S. rhizophila* species, as confirmed by a BlastP. Although the genetic cluster is highly conserved, its genomic position varies among strains, indicative of independent acquisition events. Strikingly, the downstream immunity-encoding region is highly variable despite carrying homologous genes. This observation indicated that the poly-immunity repertoire undergoes rapid evolution through gene duplications or gain and loss.

**Fig. 2.**
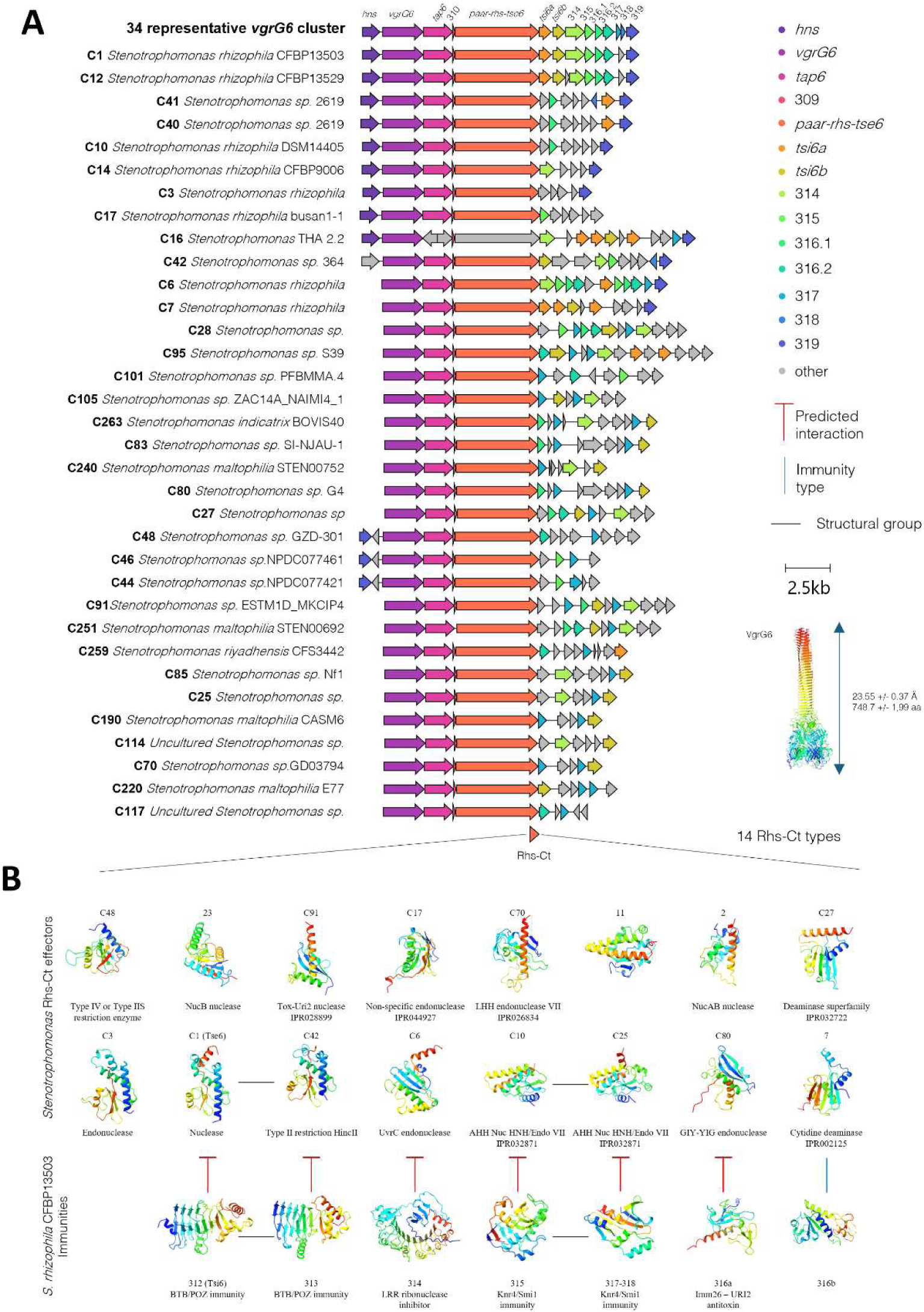
Synteny of *vgrG6* cluster and structural diversity of Rhs-Ct among the *Stenotrophomonas* genus. **A)** Conservation of *vgrG6* cluster organization in 34 *Stenotrophomonas* strains. The 14 genes from CFBP13503 *vgrG6* cluster (coloured blocks) were used as a query for a blast using Clinker online tool (CAGECAT). Thirty-four clusters were selected based on the genetic organization diversity to be shown on the figure. Each strain is indicated by its name and associated with a unique cluster number (C number). Genetic clusters are shown from *hns* to *HKJBHOBG_00319* if present. **B)** Rhs-Ct structural diversity. A blast search was performed using Clinker with PAAR-Rhs-Tse6 as a query and resulted in detecting 182 unique amino-acid sequences across 158 *Stenotrophomonas* genomes. Alignment of all sequences resulted in sorting 25 representative sequences (identified as a group or a cluster) (Fig. S3). Rhs-Ct extraction led to the identification of 14 Rhs-Ct types based on structural comparisons (Fig. S4). A putative function was assigned to each Rhs-Ct type using bioinformatic tools if possible. A vertical black bar indicates structural homologues. Horizontal red crossed bars indicate a predicted interaction between the effector and the immunity. A vertical blue bar indicates the effector-immunity pair family. The name above the structure is referring to the genetic cluster from which it is extracted.

### Structural comparisons of the Rhs-Ct revealed 14 different effectors in *Stenotrophomonas* diversity

Since Rhs are known to recombine the C-terminal domain, we compared all the *Stenotrophomonas* spp. Rhs-Ct. Based on the PAAR-Rhs-Tse6_CFBP13503_, the BlastP resulted in 182 unique amino-acid sequences detected across 158 genomes (**Fig. S3**). Subsequent sequence alignment, structure prediction and structural clustering resulted in 14 Rhs-Ct structural groups (**Fig. 2B, Fig. S4**). Functional annotation identified 18 effectors with different putative functions of the nuclease superfamily such as URI-2 nucleases, AHH nucleases, LHH RNase, GIY-YIG endonucleases, non-specific endonucleases, deaminase and a cytidine deaminase activity (**Table 1**). These putative functions were supported by the prediction of domains from the cognate immunity detected by protein-protein interaction modelling, namely Imm12, DUF4303, LRR, Knr4/Smi1, SUKH-4 and Imm1-domain containing proteins, respectively. Groups C70 (LHH nuclease), C80 (URI2-like nuclease) and C10 (AHH nuclease) are the most prevalent across *Stenotrophomonas* (**Fig. S5**). Inversely, groups 11 and C3 are the rarest while C1 (Tse6) is shared by only 2 % of the 163 *vgrG6*-positive strains. CFBP13503 carries 5 different types of immunities, of which we predicted a positive interaction for each but 316b with at least one Rhs-Ct from the *Stenotrophomonas* genus. Indeed, the putative BTB/POZ domain-containing immunity Tsi6b is predicted to block C42 endonuclease activity, 314 to block C6 UvrC-like endonucleases, 315 or 317-318 to block C10 and C25 AHH nucleases, 316a to block C80 URI2 endonucleases and 316b to block group 7 tRNA deaminases (**Table S1**). The presence of Tsi6 homologues in the diversity also suggests that *Stenotrophomonas* strains are resistant to Tse6.

**Table 1.**
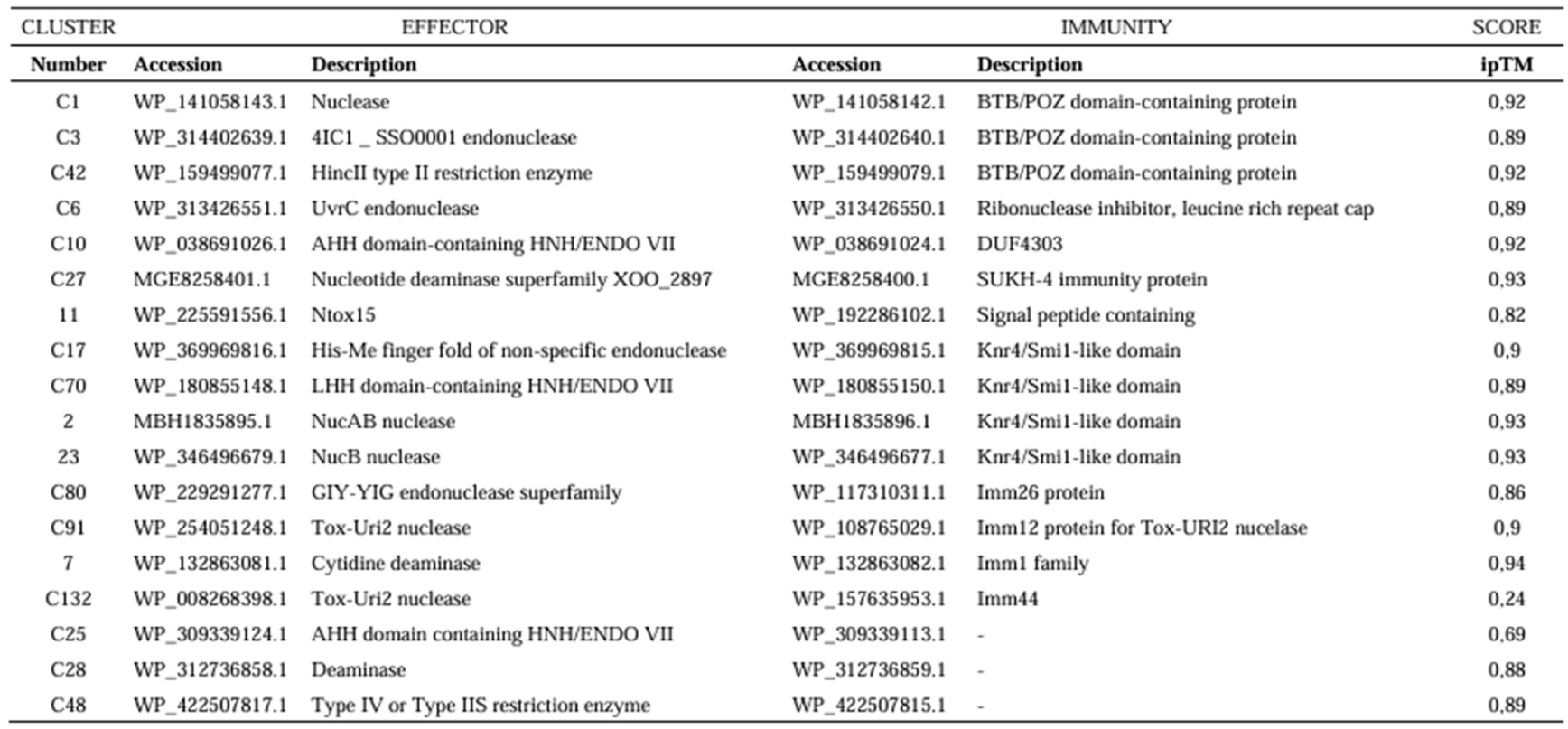
Description of the 18 Rhs-Ct types and their cognate immunity from *Stenotrophomonas* spp. Interaction prediction between the effector and the corresponding immunity was submitted to AlphaFold3 modelling resulting in the ipTM score indicated in the last column. A confident interaction is predicted by an ipTM score > 0.75.

### Tse6 is found in saline-associated Gram-negative Planctinomycetes and Gram-positive *Actinomycetes*.

The extensive diversification of Rhs-Ct domains suggests that each effector is specifically adapted to a particular ecological niche or target organism. To identify the potential environment or target relevant for Tse6_CFBP13503_ use, we searched for orthologs combining both an amino acid sequence-based blast using the NCBI database and a structure-based comparison with Foldseek server (38). This approach identified Tse6 orthologs in 37 unique strains (**Table S2**), of which 43% belongs to Gram-positive *Actinomycetes*, 22% to *γ-Proteobacteria*, 11% to *Planctomycetes*, 8% to *δ-Proteobacteria* and 5% to *α-Proteobacteria* (**Fig. 3A**). Interestingly, all *Actinomycetes* and *Planctinomycetes* strains identified are halophilic and associated with saline or aquatic environments. *S. rhyiadensis* is one of the only *Stenotrophomonas* species carrying a Tse6 homolog and was isolated from desert-associated dried and saline sand.

**Fig. 3.**
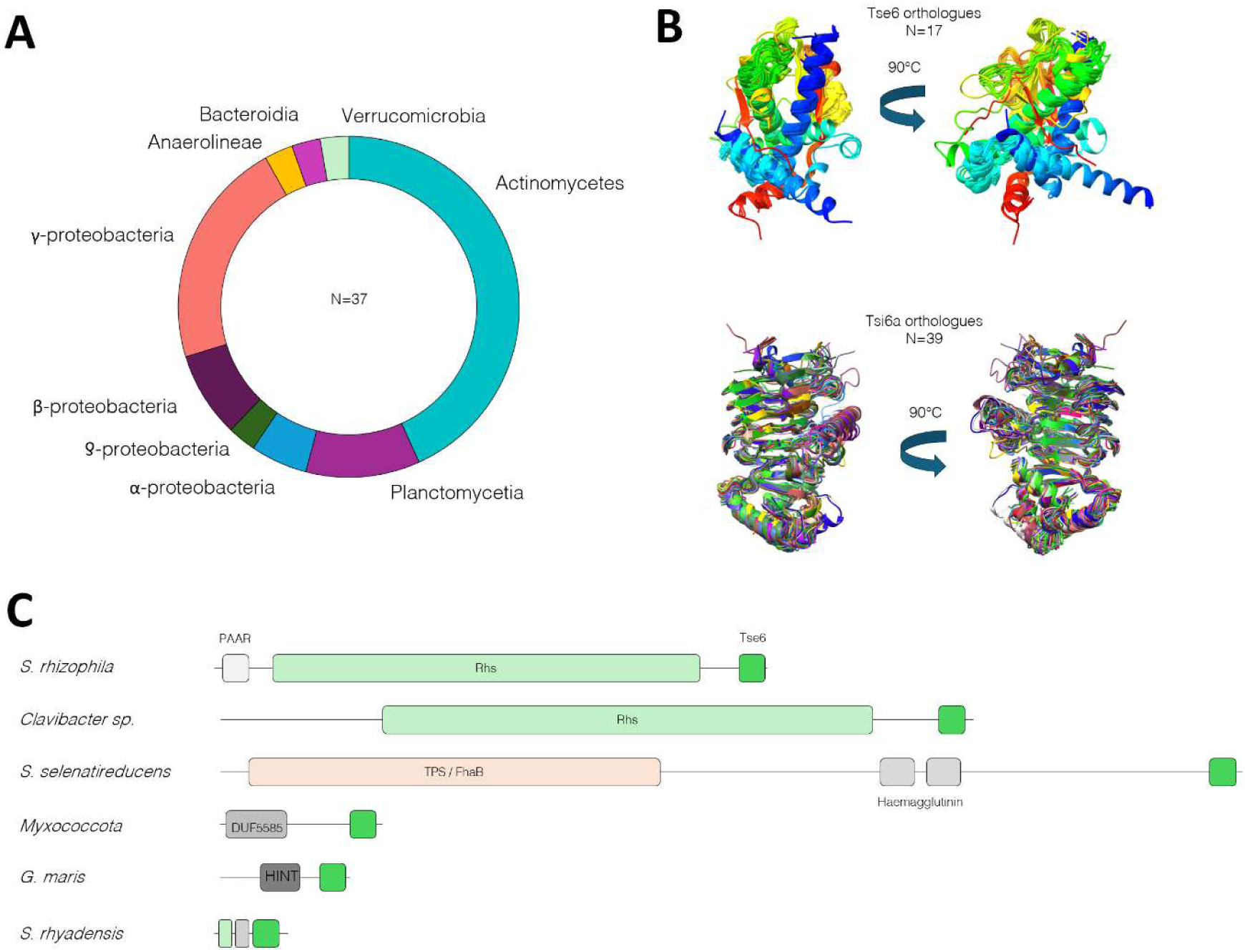
Distribution of Tse6 orthologues in bacterial diversity. **A)** Pie chart of Tse6 distribution in the bacterial diversity. Two analyses were merged to produce the pie chart composed of 37 strains. First, the amino-acid sequence of Tse6 was subjected to BlastP (NCBI database). Second, Tse6 predicted structure was subjected to a structural Blast using Foldseek and hits from AFDB1000 with *p* <0.1 were extracted. **B)** Predicted structures of Tse6 and Tsi6a orthologues. Unique Tse6 (n=17) and Tsi6a (n=39) orthologues were subjected to structure prediction by AlphaFold3 and aligned to Tse6_CFBP13503_ or Tsi6a_CFBP13503_ with ChimeraX. Structures are coloured from N-terminus (blue) to C-terminus (red) domain. **C)** Illustration of secretion signal domains found in full-length Tse6 orthologues (from top to bottom: *Stenotrophomonas rhizophila*, *Clavibacter*, *Sedimenticola selenatireducens*, *Myxococcota* bacterium, *Gimesia maris*, *Stenotrophomonas rhyadensis*).

To confirm the orthology of these hits, Tse6 and Tsi6 orthologous structures were predicted and aligned with Tse6_CFBP13503_ and Tsi6a_CFBP13503_, respectively, which we observed to perfectly match (**Fig. 3B**). Tse6 orthologues were associated with different secretion systems as indicated by the presence of PAAR, Rhs, HINT, TPS, Ig-like, lipoprotein or no specific domains in the protein sequence (**Fig. 3C**) (17). *Stenotrophomonas* strains as well as *Verrucomicrobium*, *Clavibacter* and *Curtobacterium* also associated Tse6 as a Rhs-fused effector. Intriguingly, *S. riyadhensis* possesses a Tse6 fused with a short Rhs domain, suggesting that this effector is not secreted but is available to adapt to another full-length Rhs by homologous recombination (22). Finally, Tsi6a_CFBP13503_ and Tsi6b_CFBP13503_ immunities were predicted to interact with several Tse6 orthologs (**Table S3**). Taken together, these results suggest that the Tse6-Tsi6 pair distribution is restricted to saline environments and mediates interbacterial competition between phylogenetically distant species sharing this niche.

### Effect of saline conditions on *S. rhizophila* growth and T6SS activity

We hypothesized that the Tse6 contributes to the *S. rhizophila* CFBP13503 adaptation to saline environments. To assess the impact of NaCl exposure to *S. rhizophila* physiology, we first compared growth in NaCl 1% medium against standard TSB10 medium (NaCl 0.05%). Consistent with the known halotolerance of *Stenotrophomonas* spp. strains, CFBP13503 grew significantly better under saline conditions, as indicated by a greater area under the curve and a higher final biomass in NaCl 1% medium (**Fig. 4B**). We next examined the effect of NaCl on the T6SS activity by microscopy using the fluorescent reporter strain B-GFP, in which the contractile sheath component TssB is fused to the sfGFP (26). When exposed to NaCl 1%, the CFBP13503 population exhibited almost three-fold higher T6SS+ cell frequency compared to a TSB10 culture (**Fig. 4C**). Moreover, the T6SS firing activity increased 1.5-fold in presence of NaCl 1% compared to TSB10 (**Fig. 4D**). We further measured that this NaCl-induced activity does not originate from transcriptional upregulation as several T6SS transcriptional reporters remained stable in this condition (**Fig. 4D**). Finally, the T6SS activity was also two-fold higher in standard TSB100 (NaCl 0,5%) compared to the ten-fold diluted TSB10 (NaCl 0,05%) medium, indicating the nutrient-rich condition is comparable to the NaCl 1% exposure (**Fig. S6**). This observation suggests that *S. rhizophila* and its T6SS use is adapted to saline conditions.

**Fig. 4.**
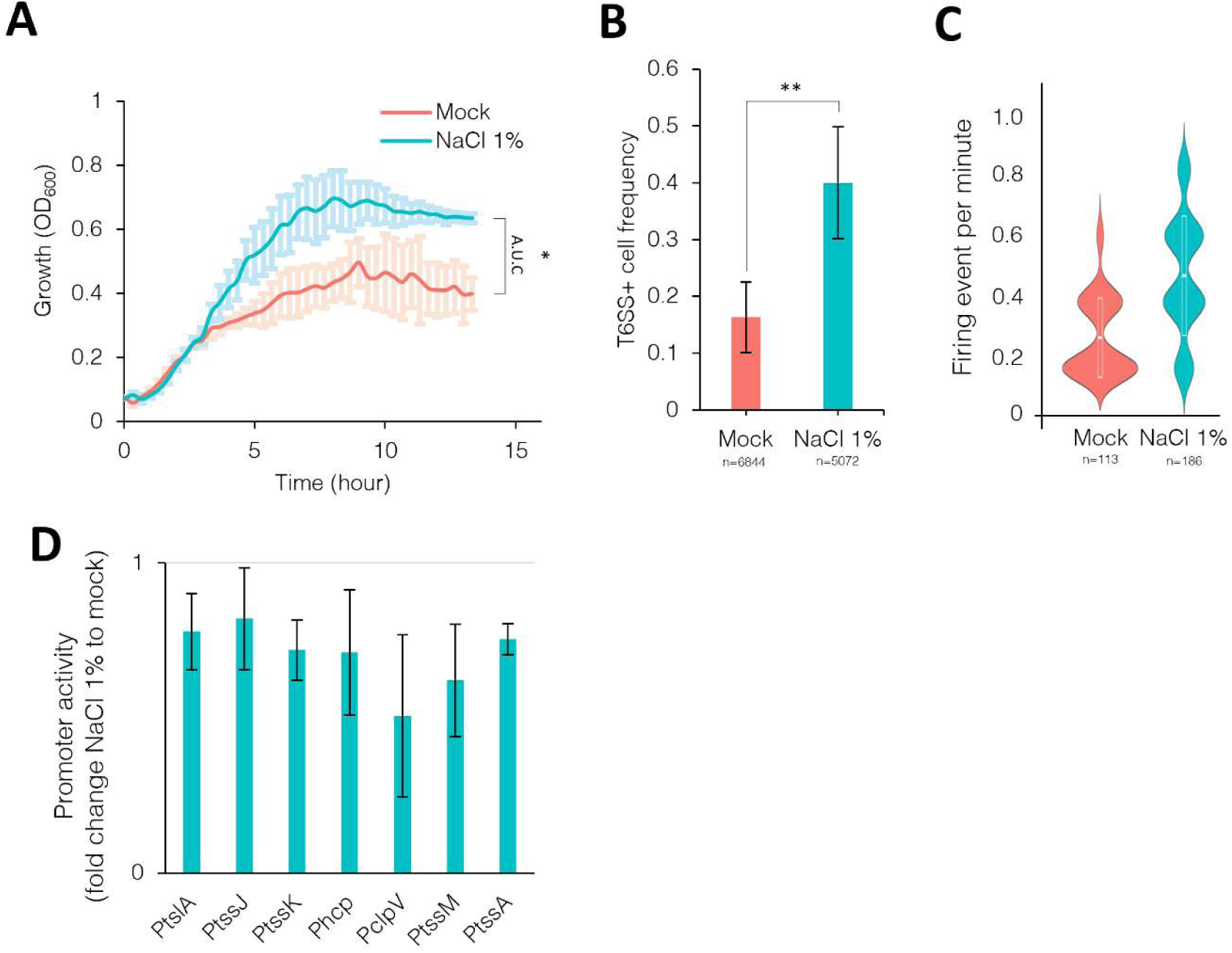
Effect of saline conditions on *S. rhizophila* CFBP13503 growth and T6SS activity. **A)** Growth kinetic of *S. rhizophila* CFBP13503 in TSB10 (Mock) or TSB10 supplemented with 1% NaCl. Data is the meaning of three biological replicates. Statistical analysis compared the aera under the curve (A.U.C) of the two conditions (*, Wilcoxon *p* <0.05). **B)** T6SS+ cell frequency in *S. rhizophila* CFBP13503 population during exponential growth in TSB10 (mock) or TSB10 supplemented with 1% NaCl. The number of cells with a T6SS focus was manually counted and the T6SS+ frequency was measured by dividing T6SS+ cell to the total cell number. Data are the means of 3 independent biological replicas with at least 3 different fields for each experiment representing 6844 (Mock) and 5072 (NaCl 1%) cells analyzed. *, Wilcoxon *p* <0.05. **C)** T6SS dynamic in *S. rhizophila* CFBP13503 population during exponential growth in TSB10 (mock) or TSB10 supplemented with 1% NaCl. Each T6SS focus was manually counted as one firing event in a 5 min time-lapse with 20 s intervals. Data combines two independent biological replicates analyzing 113 and 186 individual cells for Mock or NaCl 1% conditions, respectively. *, Wilcoxon *p* <0.05. **D)** Promoter activity of *Sr*-T6SS structural genes under NaCl exposure. Promoter activity level of structural genes in *S. rhizophila* carrying a fluorescent transcriptional reporter plasmid when grown in TSB10 supplemented with NaCl 1% compared to TSB10 (mock). Data are the mean ratio between NaCl 1% and mock conditions of at least three biological replicates. **E)** T6SS+ cell frequency in *S. rhizophila* CFBP13503 population during exponential growth in TSB10 or TSB100. The number of cells with a T6SS focus was manually counted and the T6SS+ frequency was measured by dividing T6SS+ cell to the total cell number. Data are the means of 3 independent biological replicas with at least 3 different fields for each experiment representing 755 (TSB10) and 793 (TSB100) cells analyzed. *, Wilcoxon *p* <0.05.

### Tse6 specifically targets *Actinomycetes* genera *Clavibacter* and *Curtobacterium* in saline conditions

To investigate the Gram-positive targeting specificity of Tse6, we selected a panel of bacterial strains frequently isolated from plants and seeds (25, 39), with a focus on Gram-positive bacteria. Pairwise competition assays of the selected targets against CFBP13503 WT or Δ*tse6* on TSA100 medium showed a better survival of *Plantibacter flavus* CFBP13513, *Clavibacter michiganensis* and *Curtobacterium herbarum* when facing Δ*tse6* (**Fig. 5**). It is noteworthy that these three strains were not highly susceptible to *Sr*-T6SS (∼1 log difference) when measuring the target survival against the WT compared to T6SS null mutant Δ*hcp* (**Fig. S7**), yet Tse6 deletion alone was almost sufficient to alleviate the residual susceptibility. In contrast, most Gram-positive strains were resistant to *Sr*-T6SS (**Fig. S7**) and ultimately showed no susceptibility to Tse6.

**Fig. 5.**
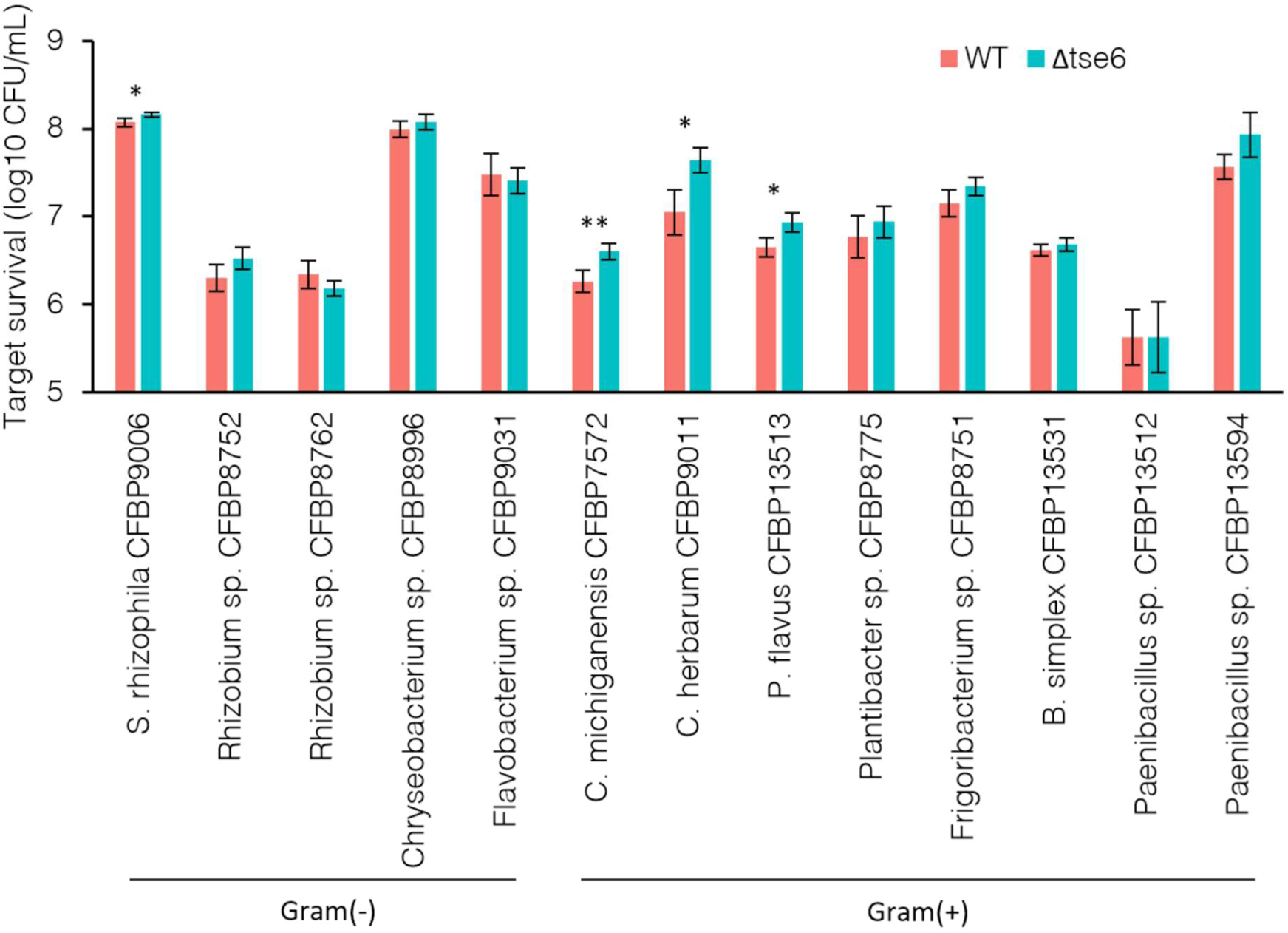
Impact of Tse6_CFBP13503_ during competition between *S. rhizophila* CFBP13503 and plant or seed-associated bacteria. Competition assay between strains isolated from plant seeds and *S. rhizophila* CFBP13503 WT or *tse6* deletion mutant. Target strain survival during 6h of coincubation on TSA100 or TSA10 for Gram-positive or Gram-negative bacteria, respectively. Data are the mean of three to four biological replicates and were subjected to a T-test (*p*<0.05).

## DISCUSSION

A comprehensive analysis of the *vgrG6* cluster of *Stenotrophomonas rhizophila* CFBP13503 reveals a highly specialized and adaptable type VI secretion system (T6SS). The genetic organization of this cluster is built around a unique PAAR-Rhs-Tse6 effector associated with a set of cognate and non-cognate immunities, forming a versatile poly-immunity module. One striking feature of this cluster is the unusually long VgrG6 tip compared to other VgrGs in *S. rhizophila* CFBP13503 (40). Structural comparisons revealed that VgrG6 length is similar to the anti-Gram-positive RhsB-associated VgrG3 (34) as well as the anti-fungal VgrG5 (35)from *Paracidovorax citrulli* but not to the anti-Gram-positive VgrGs from *Acinetobacter baumannii* (41) and *Aeromonas dhakensis* (42), which display shorter tip (**Fig. S2**). *P. citrulli* VgrG3-associated effector is thought to act in the cytoplasm whereas effectors from *A. baumannii* and *A. dhakensis* display anti-bacterial activity from the extracellular milieu, likely explaining the need for a long and a short tip, respectively. Although *S. rhizophila* and *P. citrulli* share similar VgrG length, the predicted structures of their associated effector Tse6 and RhsB-Ct substantially differ, suggesting a distinct mode of action.

Comparative genomics demonstrates that the *vgrG6* cluster core genes (*vgrG6*, *tap6*, *paar-rhs-tse*) are broadly conserved across *Stenotrophomonas* species, with homologous clusters detected in 164 out of 193 genomes, underscoring the evolutionary importance of this feature within the genus (**Fig. 2**). Inversely, the downstream poly-immunity region is highly variable across strains, reflecting a key role for this cluster in the strain adaptation. Importantly, this region concerns a poly-immunity cluster. This architecture suggests a strong selective pressure linked to bacterial interactions and an extended protection mechanism against the diversity of effectors present in potential competitors. It has recently been reported that poly-immunity clusters work as a direct protection against threats from the cognate and non-cognate effectors in microbial competition, notably in the microbiota context (7–9, 43). Due to the high selective pressure imposed by effector exposition, gain and loss of effector-immunity pairs likely reflects an arm race as an adaptive process to ecological niches (11, 12, 22, 44).

Interbacterial genetic exchanges represent a major driver of strain diversification and adaptation. Numerous comparative genomic studies have documented the direct acquisition of effector-immunity pairs by horizontal gene transfer (11, 12). Other studies identified genetic determinants for gene exchange and recombination such as the presence of poly-immunity clusters on mobile genetic elements in *Bacteroides* (7) or insertion sequence sites and Xer-dependent recombination systems next to poly-immunities in *Xenorhabdus* and *Photorhabdus* (6). In *Stenotrophomonas*, we similarly observed some *vgrG6* clusters flanked by integrases and tRNA coding genes, indicative of independent horizontal acquisition events, further supported by atypical GC content.

Systematic analysis of the *vgrG6* cluster-associated PAAR-Rhs protein found in 163 *Stenotrophomonas* genomes revealed that this protein is prone to evolution on both the core and the C-terminal domains. First, PAAR-less Rhs or size-variable Rhs domains can be found, indicative of high selective pressure even on the core Rhs domain. Second, structural analyses of the Rhs-Ct extracted revealed a wide diversity of effectors, once again highlighting the extensive plasticity of Rhs proteins. These Rhs-Ct grouped into 14 distinct families (structural cluster) (**Fig. 2**), among which we identified putative deaminase and nuclease activities such as putative GIY-YIG endonucleases, AHH, LHH, Tox-Uri2, Ntox15 and NucB-containing domain nucleases. These nuclease-related functions are consistent with a cytoplasmic effector, usually found associated with Rhs as a polymorphic toxin that acts on DNA, RNA or related molecules such as tRNA, NADH or ribosomes (6, 17). For each Rhs-Ct family, we identified the putative functional domain of the cognate immunity. These immunities were identified as BTB/POZ, DUF4303, Knr4/Smi1, SUKH-4, Imm and LRR domain-containing proteins, consistent with the putative function of the cognate effector. Taken together, these findings confirmed that the Rhs-Ct and associated immunities are highly variable and quickly exchangeable in the *Stenotrophomonas* genus diversity, reflecting a high adaptive potential for this PAAR-Rhs effector.

It is important to note that our previous proteomic analysis detected the production of three *vgrG6*-associated immunities in *S. rhizophila* CFBP13503, of which Tsi6a, Tsi6b and the LRR domain-containing protein 314 (40). This underlies the importance for cross-protection against Tse6 orthologs and LRR cognate effectors in *S. rhizophila* CFBP13053. One could hypothesize that the other non-expressed immunities could be regulated under specific conditions, as suggested by the detection of putative promoters for 315, 316a and 317 genes. Interestingly, LRR domain-containing proteins are described as involved in plant immunity. They notably recognize diverse pathogen-associated molecular pattern proteins (e.g. T3SS effectors) that ultimately triggers their function for immunity pathway activation, such as signalisation, gene regulation or pore-forming activity (45, 46). In *Burkholderia gladioli*, nuclease immunities have been found to carry HTH DNA binding domain thus playing a role in gene regulation upon effector neutralization (47). Future work should investigate whether *vgrG6* cluster immunities are inducible and involved in a more global defense program. In conclusion the *vgrG6*-associated immunities expand the defensive repertoire through diversification and cross-protection.

Throughout this study, effector-immunity interaction predictions were performed exclusively using AlphaFold3 (48), which has demonstrated high precision in predicting both protein structures and protein-protein interactions applied to effector-immunity studies. This approach is well supported by recent studies confirming that AlphaFold-based predictions reliably capture the effector-immunity biochemistry that has been confirmed *in vivo* (6, 21, 27, 28). However, our findings on the immunity cross-protection should be confirmed by toxicity assays in the future.

Rhs-Ct domains belonging to the same structural family sometimes occur in markedly distinct genomic environments (e. g. Cluster 95 and 48 or 220 and 42), consistent with independent acquisition events occurring at multiple evolutionary timescales during adaptive process in *Stenotrophomonas*. Conversely, related Rhs-Ct families can display a highly divergent amino acid sequence, suggesting an acquisition from a different organism or environment. Together, these observations support a model in which Rhs-Ct effectors are repeatedly and independently acquired from the microbial community to adapt competitive interactions with local competitors. More importantly, Rhs-Ct types are differently distributed across bacterial genera, some are broadly shared while others, such as Tse6, appear much rarer. One could hypothesize that such a rarity reflects a high degree of specificity. Indeed, ortholog searches based on structural similarities and amino-acid sequence identified *Actinomycetes* and *Planctinomycetes* as the dominant taxonomic groups out of 37 Tse6-carrying strains (**Fig. 3**). The γ-Proteobacteria class, of which *S. rhizophila* belongs, is not the majority among these hits, it suggests that CFBP13503 deploys Tse6 in a highly specific ecological niche dominated by Gram-positive *Actinomycetes* and Gram-negative *Planctomycetes*. This conclusion is further supported by the predicted interaction of Tsi6a_CFBP13503_ and Tsi6b_CFBP13503_ immunities with several Tse6 orthologs from the identified *Planctomycetes* strains. Using the phytopathogen *Clavibacter michiganensis* CFBP7572 (49) and the plant beneficial bacteria *Curtobacterium herbarum* CFBP9011 (50, 51) and *Plantibacter flavus* CFBP13531, we confirmed the activity of Tse6 against *Actinomycetes*, though only the latter was co-isolated with *S. rhizophila* CFBP13503 (25). Since Gram-negative bacteria were not susceptible to Tse6, we finally confirmed the role for Tse6 in targeting Gram-positive strains and ultimately correlating the functional link between VgrG6 length and such a target (40). We do not exclude a target specificity towards fungi, still to be explored.

One limitation of our ortholog search is the potential influence of codon usage bias: a BlastP query using Tse6 from CFBP13503 returned only closely related sequences, likely underestimating true ortholog diversity. Using instead the *Clavibacter*-derived Tse6 ortholog WP_194665312 as a query revealed additional, previously undetected orthologs in *Streptomyces* and *Rathayibacter* strains, suggesting that the true taxonomic distribution of this effector family extends further than initially identified.

The presence of multiple Rhs-Ct immunity types within *vgrG6* clusters across *Stenotrophomonas* strains suggests that poly-immunity cluster serves as a broad-spectrum defense against intraspecific competitors. Consistent with this, the CFBP13503 *vgrG6* cluster encodes seven immunity genes, of which all but 316b are predicted to neutralize a specific Rhs-Ct family. This observation leads us to speculate that *Stenotrophomonas* strains use the *vgrG6* cluster as an offensive and defensive system for intra-specific competition. The concept of kin discrimination has been discussed in recent studies to explain the extensive diversification of polymorphic toxins at the genus and species levels (33, 42). As evidence, strains from the same species evolve new toxin modules while collecting immunities from ancestral ones, thereby gaining competitive advantages over close relatives (7, 9, 52–55). However, this model does not apply to Tse6, which we showed to not be involved in competition with phylogenetically close bacteria, including susceptible *S. rhizophila* isolated from similar niches (40). Consistently, deletion of *tse6* had no significant effect on the survival of *S. rhizophila* CFBP9006 (**Fig. 5**), a strain lacking both the *vgrG6* cluster and *tsi6* immunities (40). Moreover, structural blasts revealed that the Rhs-Ct effectors identified in this study are broadly distributed across bacterial diversity, suggesting that the *vgrG6* cluster equips *S. rhizophila* to compete with, and defend against, a wide range of phylogenetically distant species rather than close relatives. By contrast, the kin discrimination should be stated when identifying a specific effector selectively targeting a closely related strain, as illustrated in *Vibrio* strains with the TseV effector (56). Effectors with such narrow phylogenetic specificity have also been identified in the T6SS repertoire of *S. rhizophila* CFBP13503, notably amidases with potent activity against bacteria from the same family (40).

Integrating these observations, and consistent with proposed models for poly-immunity roles (52), we propose the following adaptive model for the *vgrG6* cluster in *Stenotrophomonas*: the strain accumulates orthologous immunities from cognate effectors of niche-invading competitors; the strain that is prone to effector-immunity diversification increased by Rhs plasticity and then becomes a new competitor of the ancestral strain; this finally leads to enhanced fitness enabling for new niche colonization by targeting new bacterial targets (**Fig. 6**). Therefore, the *vgrG6* system balances kin discrimination and broad-spectrum antagonism in bacterial communities. In our study, we applied a good methodology to challenge this model since competition assays were performed between strains co-isolated from the same plant seeds and seedlings habitat (25).

**Fig. 6.**
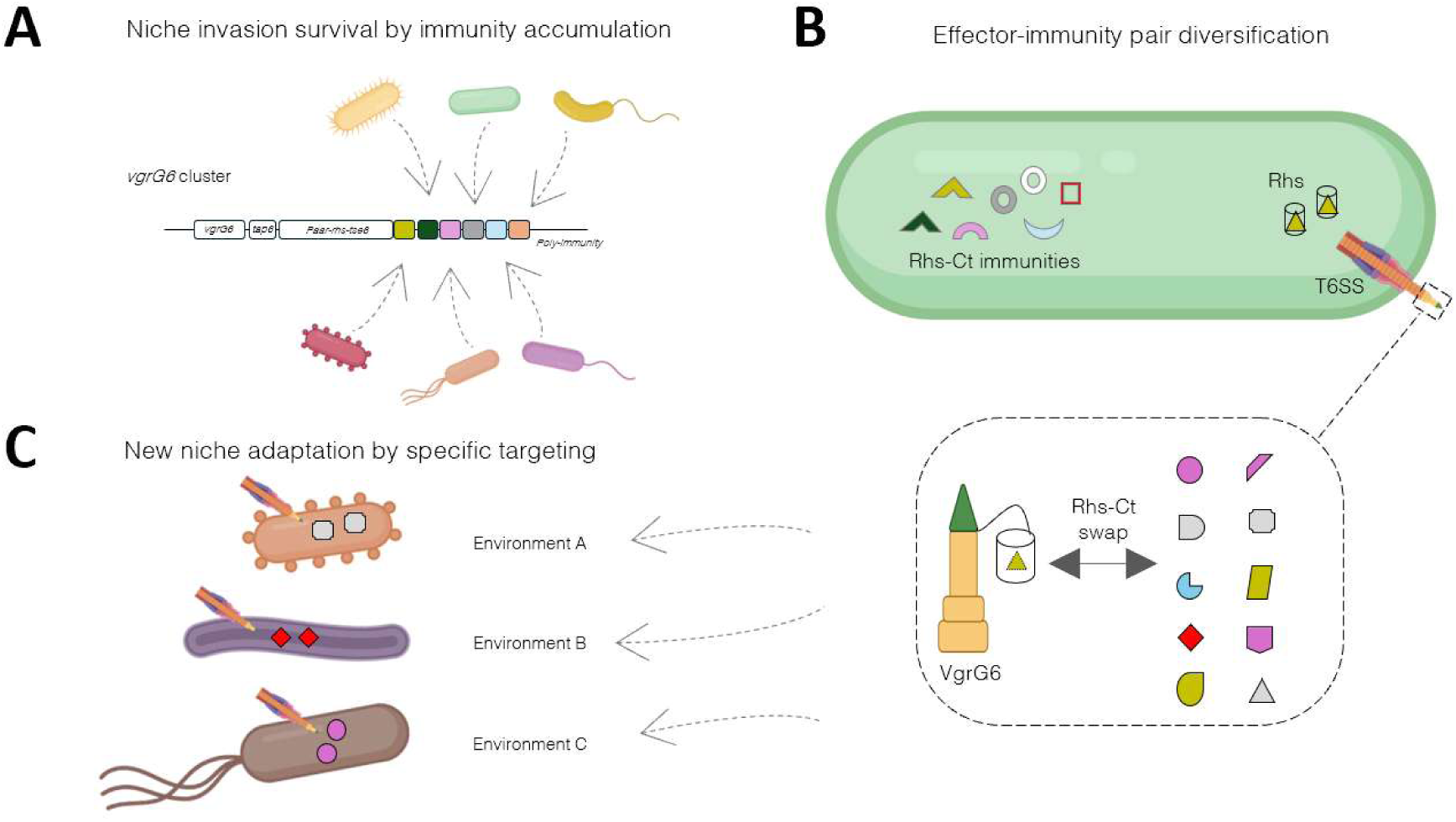
Model of vgrG6 cluster adaptation and mode of action. The *vgrG6* cluster from *Stenotrophomonas* strains accumulates orthologous immunities from cognate effectors of niche-invading competitors (**A**). The strain is prone to effector-immunity diversification through Rhs plasticity and then becomes a new competitor of the ancestral strain (**B**). This finally leads to enhanced fitness, enabling for new niche colonization by targeting new bacterial targets (**C**).

We finally investigated the effect of saline conditions on *Sr*-T6SS activity (**Fig. 4**). *S. rhizophila* is known to play a role in salt stress tolerance of plants and is naturally adapted to saline environments (57), consistent with results obtained in the presented study. *S. rhizophila* DSM14405 displays antifungal activity (58) and T6SS genes are upregulated (59, 60) under high salt conditions. However, a higher T6SS activity in NaCl 1% was not accompanied by T6SS gene upregulation in CFBP13503, suggesting a post-translational regulation. Interestingly, the genomic environment of the *vgrG6*_CFBP13503_ cluster is composed of a *degQ* homolog, a periplasmic pH-dependent endoprotease that could respond to saline stress (61). Furthermore, Tse6 orthologues were mostly found with saline-adapted bacteria such as *Actinomycetes* and *Planctomycetes* (**Table S2**). Then, future work should determine how NaCl modulates *Sr*-T6SS regulation and whether *degQ* system is involved.

Taken together, our findings support a model in which Tse6 is deployed in a specific environment where *S. rhizophila* CFBP13503 likely competes with a saline-adapted microbial community. This study demonstrates that the Rhs-containing *vgrG6* cluster confers a conserved, high adaptive potential in *Stenotrophomonas* species through defensive and offensive strategy. It supports a model in which effector innovation and immunity diversification are key drivers of ecological specialization.

## MATERIAL AND METHOD

### Bacterial strains and growth conditions

Strains used in this study are listed in **Table S4**. Gram-positive strains were grown in tryptic soy agar (TSA100, NaCl 0.5%) or broth (TSB100). *S. rhizophila* and other Gram-negative strains were grown in TSB10 (TSB100 diluted ten-fold, NaCl 0.05%). The richest TSA100 medium limited cyst formation and dormant stage in the tested Gram-positive strains. All strains were grown in a 13 mL plastic tube and incubated at 28°C with agitation at 150 rpm. When needed, cultures were supplemented with NaCl 10 mg/mL (1%) antibiotic as follows: tetracycline 20 µg/mL (Tc20), rifamycin 50 µg/mL (Rif50), rifamycin 200 µg/mL (Rif200), or polymyxin E 25 µg/mL (PE25).

### Growth kinetic

Fitness of strains was assessed by a growth kinetic assay in microplates. Typically, overnight cultures in TSB10 or TSB100 were incubated at 28°C with agitation. Cultures were then washed and adjusted to an OD_600_ 0.5 with fresh medium supplemented or not with NaCl 1% and appropriate antibiotics. Samples were ten-fold diluted in 200 µL of the appropriate medium in a 96-flat transparent well microplate (Greiner Bio-One) and incubated in a microplate reader (VANTAstar, BMG Labtech) for 24h at 28°C with 300 rpm double orbital agitation. Growth curves were drawn from the average of three technical replicates of three biological replicates. Area under the curve was also calculated using *growthcurver* package (62) under R studio software.

### Spontaneous rifamycin resistance

For selective recovery of target strains, spontaneous rifamycin-resistance was produced by plating 100 µL of a concentrated exponential culture on TSA100+Rif200. Recovered colonies (RifR) were spread again on TSA100+Rif200. Fitness of RifR clones was tested by comparing growth with the wild-type strain in a growth kinetic assay in both TSB100 and TSB100+Rif50. The fittest clones were then used to check for a possible reversion by growing them in TSB100+Rif50 after an overnight culture in TSB100. Recovered cultures in TSB100+Rif50 were saved in glycerol 15% at -80°C.

### Interbacterial competition assay

Strains isolated from plant seeds (target strains) were subjected to competition assay against *S. rhizophila* WT or deletion mutants Δ*hcp* and Δ*tse6* (attacker strains). To do so, Gram-positive target strains were grown overnight in TSB100 and diluted in fresh TSB100 for exponential growth prior to being mixed with attackers. Gram-negative target strains and *S. rhizophila* attackers were grown for 17h in TSB10. Target strains and attackers were prepared at OD_600_ 2 and OD_600_ 1, respectively, and mixed in an equal volume. Ten µL of the mixture was then spotted on TSA100 or TSA10 for Gram-positive or negative target strains and incubated 4h or 6h at 28°C, respectively. Spots were then resuspended in 1 mL TSB100 and serially ten-fold diluted. Twenty microliters of each dilution were spotted on TSA100+Rif50 for RifR strains or TSA100+PE25 for other strains naturally resistant to polymyxin E. Colony forming unit (CFU) of recovered target strains was then counted and transformed according to the dilution factor to express target survival in log_10_ CFU/mL.

### Fluorescent reporter assays

Fluorescence reporter strains were used to test the promoter activity of *Sr*-T6SS structural genes in saline conditions. Cells with pME transcriptional fusions or the empty vector (40) were grown in TSB10+Tc20 overnight at 28°C with agitation at 150 rpm. Overnight cultures were diluted into fresh TSB10 (mock) or TSB10 supplemented with NaCl 1% and grown for 5h at 28°C with agitation. Cultures were then adjusted to OD_600_ 1 and 200 μL was dispensed into a 96-well black clear-bottom microplate (Greiner Bio-One). Fluorescence of the mCherry (F_570_, excitation: 520 nm, emission: 570 nm) and sample absorbance (OD_600_) were measured using the VANTAstar microplate reader (BMG Labtech). Fluorescence values were normalized to the corresponding OD_600_ values using the formula F_570_/OD_600_. Normalized fluorescence values were subtracted from the empty vector values and a ratio between NaCl and mock conditions was calculated. Results are thus expressed in fluorescence fold change for NaCl conditions relative to mock.

### Comparative genomics

Comparative genomics of the CFBP13503 *vgrG6* cluster was performed with the online tools Clinker and Cblaster from CAGECAT (37). Basically, the genetic region encompassing the *vgrG6* cluster as described in **Fig. 1** was subjected to a Clinker blast within *Stenotrophomonas* genus using default mode (job P470W541W183W19). The analysis reported 236 genetic clusters across 164 bacterial genomes. We aligned the first 50 genetic clusters and selected 34, illustrating the diversity that can be found in *Stenotrophomonas* spp. in **Fig. 2**. Then, only PAAR-Rhs-Tse6 (WP_141058143.1) was subjected to Cblaster research in *Stenotrophomonas* genus with default mode, resulting in 255 hits from 163 bacterial genomes (job X102A714F137F00). All homologous PAAR-Rhs sequences were extracted and aligned to draw a phylogenetic tree (**Fig. S3**) using Clustal Omega (EMBL-EBI) and therefore defining 25 different types. The 25-representative C-terminal domains (Rhs-Ct) from these sequences were isolated and subjected to functional analysis using InterProScan (63) and structural prediction using AlphaFold3 (48). Structure comparisons by DALI dendrograms (64) resulted in defining 14 structural groups (**Fig. S4**). Each Rhs-Ct predicted structure was subjected to Foldseek blasting (38) to measure its distribution (**Fig. S5**). This analysis detected Rhs-Ct groups in 4,583 unique bacterial strains from 1,308 unique genera (e-value <1). Finally, cognate immunities for each associated Rhs-Ct were extracted from representative clusters and effector-immunity pairs were tested using AlphaFold3 (positive interaction for ipTM > 0.75). *vgrG6*_CFB13503_ cluster-associated immunities were also tested for their potential interaction (i.e. inhibition) with other Rhs-Ct types.

Tse6 (Rhs-Ct_CFBP13503_) amino-acid sequence or predicted structure was subjected to BlastP (NCBI) or Foldseek analysis, respectively. The first allowed for finding of closest orthologous sequences while the latter allowed for finding the best structural orthologues. BlastP sequences were extracted and modelled with AlphaFold3 to check the structural similarity with Tse6. Sequences were also subjected to InterProScan to identify a potential domain indicative of the secretion mode (PAAR, Rhs, HINT, Ig-like, lipoprotein, TPS). Tsi6a and Tsi6b were tested for a potential interaction with Tse6 orthologues using AlphaFold3 (ipTM > 0.75).

### Fluorescence microscopy

The *Sr*-T6SS activity was measured by fluorescence microscopy. We used *S. rhizophila* B-GFP strain in which the major T6SS sheath subunit *tssB* is tagged with the sfGFP, allowing for visualization and tracking of the T6SS in the cell (40). The microscope used was a Zeiss Axio Imager Z2 equipped with an Axiocam 305 color camera, at iMAC (IRHS cellular imaging platform, SFR4207 QuaSaV). The filters were composed of a 470/40 nm bandpass excitation filter, a 495 nm beamsplitter, and a 525/50 nm bandpass emission filter for GFP fluorescence capture with 20% light and 100 ms exposure time. Typically, B-GFP was cultured overnight in TSB10 and diluted for exponential growth in TSB10 or in TSB10 supplemented with NaCl 1% for 5h at 28°C. Cells were then centrifuged and concentrated ten-fold. One microliter of the sample was spotted onto a microscope slide poured with a 2% agarose pad (NuSieve, BMA) in a gene frame and covered with a coverslip. T6SS presence was defined by a GFP focus in the cell, and T6SS+ cell frequency was calculated as the number of cells with a T6SS out of the total number of cells. Measures were performed on at least 3 different fields for each biological replicates and image analysis was performed with Fiji (ImageJ) software (65).

## ACKNOWLEDGMENT

We thank Chrystelle Brin, Audrey Latus, and Marion Barbier for technical assistance, the IMAC platform for imaging facilities (SFR4207 QuaSaV), Martial Briand for support in genomics and the CIRM-CFBP for providing access to strains from its collection (https://doi.org/10.15454/E8XX-4Z18). This work was supported by the Institut National de Recherche pour l’Agriculture, l’Alimentation et l’Environnement (INRAE), Université Angers, and Angers Loire Métropole (T6IMPACT project).

## REFERENCES

1. Cianfanelli FR, Monlezun L, Coulthurst SJ. 2016. Aim, Load, Fire: The Type VI Secretion System, a Bacterial Nanoweapon. Trends in Microbiology 24:51–62.

2. Unni R, Pintor KL, Diepold A, Unterweger D. 2022. Presence and absence of type VI secretion systems in bacteria. Microbiology 168:001151.

3. Cherrak Y, Flaugnatti N, Durand E, Journet L, Cascales E. 2019. Structure and Activity of the Type VI Secretion System. Microbiology Spectrum 7: 10.1128/microbiolspec.psib-0031–2019.

4. Jurėnas D, Journet L. 2021. Activity, delivery, and diversity of Type VI secretion effectors. Molecular Microbiology 115:383–394.

5. Hagan M, Pankov G, Gallegos-Monterrosa R, Williams DJ, Earl C, Buchanan G, Hunter WN, Coulthurst SJ. 2023. Rhs NADase effectors and their immunity proteins are exchangeable mediators of inter-bacterial competition in Serratia. Nat Commun 14:6061.

6. Martinkus J, Ginibre-Supersac N, Rivard N, Belot L, Brillard J, Cascales E, Jurėnas D. 2026. Poly-immunity arrays associated with Rhs toxins confer wide protection against competitors. Current Biology 36:176–187.e3.

7. Ross BD, Verster AJ, Radey MC, Schmidtke DT, Pope CE, Hoffman LR, Hajjar AM, Peterson SB, Borenstein E, Mougous JD. 2019. Human gut bacteria contain acquired interbacterial defence systems. Nature 575:224–228.

8. Knecht A, Sirias D, Utter DR, Gibbs KA. 2025. Non-cognate immunity proteins provide broader defenses against interbacterial effectors in microbial communities. eLife 12:RP90607.

9. Martinkus J, Ginibre-Supersac N, Rivard N, Belot L, Brillard J, Cascales E, Jurėnas D. 2026. Poly-immunity arrays associated with Rhs toxins confer wide protection against competitors. Current Biology 36:176–187.e3.

10. Azhieh A, Hernandez P, Anderson AC, Sychantha D, Verster AJ, Whitney JC, Ross BD. 2025. Rapidly evolving orphan immunity genes protect human gut bacteria from intoxication by the type VI secretion system. bioRxiv 10.1101/2025.05.03.651265.

11. García-Bayona L, Coyne MJ, Comstock LE. 2021. Mobile Type VI secretion system loci of the gut Bacteroidales display extensive intra-ecosystem transfer, multi-species spread and geographical clustering. PLOS Genetics 17:e1009541.

12. Habich A, Chaves Vargas V, Robinson LA, Allsopp LP, Unterweger D. 2025. Distribution of the four type VI secretion systems in Pseudomonas aeruginosa and classification of their core and accessory effectors. Nat Commun 16:888.

13. Jackson AP, Thomas GH, Parkhill J, Thomson NR. 2009. Evolutionary diversification of an ancient gene family (rhs) through C-terminal displacement. BMC Genomics 10:584.

14. Zhang D, Iyer LM, Aravind L. 2011. A novel immunity system for bacterial nucleic acid degrading toxins and its recruitment in various eukaryotic and DNA viral systems. Nucleic Acids Res 39:4532–4552.

15. Ma J, Sun M, Dong W, Pan Z, Lu C, Yao H. 2017. PAAR-Rhs proteins harbor various C-terminal toxins to diversify the antibacterial pathways of type VI secretion systems. Environmental Microbiology 19:345–360.

16. Ma J, Sun M, Dong W, Pan Z, Lu C, Yao H. 2017. PAAR-Rhs proteins harbor various C-terminal toxins to diversify the antibacterial pathways of type VI secretion systems. Environmental Microbiology 19:345–360.

17. Zhang D, de Souza RF, Anantharaman V, Iyer LM, Aravind L. 2012. Polymorphic toxin systems: Comprehensive characterization of trafficking modes, processing, mechanisms of action, immunity and ecology using comparative genomics. Biol Direct 7:18.

18. Ruhe ZC, Low DA, Hayes CS. 2020. Polymorphic Toxins and Their Immunity Proteins: Diversity, Evolution, and Mechanisms of Delivery. Annual Review of Microbiology 74:497–520.

19. Li H, Tan Y, Zhang D. 2022. Genomic discovery and structural dissection of a novel type of polymorphic toxin system in gram-positive bacteria. Computational and Structural Biotechnology Journal 20:4517–4531.

20. Kobayashi K. 2021. Diverse LXG toxin and antitoxin systems specifically mediate intraspecies competition in Bacillus subtilis biofilms. PLOS Genetics 17:e1009682.

21. Wang F, Luo J, Zhang Z, Liu Y, Sheng D hong, Zhuo L, Li Y. 2025. Differential crosstalk between toxin-immunity protein homologs divides Myxococcus nonself siblings into close and distant social relatives. mBio 16:e03902–24.

22. Koskiniemi S, Garza-Sánchez F, Sandegren L, Webb JS, Braaten BA, Poole SJ, Andersson DI, Hayes CS, Low DA. 2014. Selection of Orphan Rhs Toxin Expression in Evolved Salmonella enterica Serovar Typhimurium. PLOS Genetics 10:e1004255.

23. Garin T, Brin C, Préveaux A, Brault A, Briand M, Simonin M, Barret M, Journet L, Sarniguet A. 2024. The type VI secretion system of *Stenotrophomonas rhizophila* CFBP13503 limits the transmission of *Xanthomonas campestris* pv. *campestris* 8004 from radish seeds to seedlings. Molecular Plant Pathology 25:e13412.

24. Wolf A, Fritze A, Hagemann M, Berg G. 2002. Stenotrophomonas rhizophila sp. nov., a novel plant-associated bacterium with antifungal properties. International Journal of Systematic and Evolutionary Microbiology 52:1937– 1944.

25. Garin T, Brault A, Marais C, Briand M, Préveaux A, Bonneau S, Simonin M, Barret M, Sarniguet A. 2025. T6SS-mediated competition by Stenotrophomonas rhizophila shapes seed-borne bacterial communities and seed-to-seedling transmission dynamics. mSystems 10:e00457–25.

26. Taillefer B, Dairy Y, Brault A, Briand M, Charpin P, Armengaud J, Sarniguet A. 2026. Specific and broad-spectrum antibacterial effectors of type VI secretion system drive competition of *Stenotrophomonas rhizophila* against bacteria from seed microbiota. Microbiol Spectr e03532–25.

27. Geller AM, Shalom M, Zlotkin D, Blum N, Levy A. 2024. Identification of type VI secretion system effector-immunity pairs using structural bioinformatics. Mol Syst Biol 20:702–718.

28. Danov A, Pollin I, Moon E, Ho M, Wilson BA, Papathanos PA, Kaplan T, Levy A. 2024. Identification of novel toxins associated with the extracellular contractile injection system using machine learning. Mol Syst Biol 20:859–879.

29. Waller PR, Sauer RT. 1996. Characterization of degQ and degS, Escherichia coli genes encoding homologs of the DegP protease. Journal of Bacteriology 178:1146–1153.

30. Krojer T, Garrido-Franco M, Huber R, Ehrmann M, Clausen T. 2002. Crystal structure of DegP (HtrA) reveals a new protease-chaperone machine. Nature 416:455–459.

31. Mikolčević P, Hloušek-Kasun A, Ahel I, Mikoč A. 2021. ADP-ribosylation systems in bacteria and viruses. Computational and Structural Biotechnology Journal 19:2366–2383.

32. Holberger LE, Garza-Sánchez F, Lamoureux J, Low DA, Hayes CS. 2012. A novel family of toxin/antitoxin proteins in Bacillus species. FEBS Letters 586:132–136.

33. Shu X, Sun X, Wang K, Duan Y, Liu Y, Zhang R. 2025. LXG Toxins of Bacillus Velezensis Mediate Contact-Dependent Inhibition in a T7SS-Dependent Manner to Enhance Rhizosphere Adaptability. International Journal of Molecular Sciences 26:2592.

34. Pei T-T, Kan Y, Wang Z-H, Tang M-X, Li H, Yan S, Cui Y, Zheng H-Y, Luo H, Liang X, Dong T. 2022. Delivery of an Rhs-family nuclease effector reveals direct penetration of the gram-positive cell envelope by a type VI secretion system in Acidovorax citrulli. mLife 1:66–78.

35. Yan S, Zou Y, Wu T, Kan Y, Luo H, Pei T-T, Liang X, An Y, Meng P, Song Y, Qin W-M, Chen C, Dong T. 2025. A broad-spectrum anti-fungal effector dictates bacterial-fungal interkingdom interactions. PLoS Pathog 21:e1013598.

36. Wiedenbeck J, Cohan FM. 2011. Origins of bacterial diversity through horizontal genetic transfer and adaptation to new ecological niches. FEMS Microbiol Rev 35:957–976.

37. Gilchrist CLM, Booth TJ, van Wersch B, van Grieken L, Medema MH, Chooi Y-H. 2021. cblaster: a remote search tool for rapid identification and visualization of homologous gene clusters. Bioinformatics Advances 1:vbab016.

38. van Kempen M, Kim SS, Tumescheit C, Mirdita M, Lee J, Gilchrist CLM, Söding J, Steinegger M. 2024. Fast and accurate protein structure search with Foldseek. Nat Biotechnol 42:243–246.

39. Simonin M, Briand M, Chesneau G, Rochefort A, Marais C, Sarniguet A, Barret M. 2022. Seed microbiota revealed by a large-scale meta-analysis including 50 plant species. New Phytologist 234:1448–1463.

40. Taillefer B, Dairy Y, Brault A, Briand M, Charpin P, Armengaud J, Sarniguet A. 2026. Specific and broad-spectrum antibacterial effectors of type VI secretion system drive competition of Stenotrophomonas rhizophila against bacteria from seed microbiota. Microbiology Spectrum 0:e03532–25.

41. Le N-H, Peters K, Espaillat A, Sheldon JR, Gray J, Di Venanzio G, Lopez J, Djahanschiri B, Mueller EA, Hennon SW, Levin PA, Ebersberger I, Skaar EP, Cava F, Vollmer W, Feldman MF. 2020. Peptidoglycan editing provides immunity to Acinetobacter baumannii during bacterial warfare. Science Advances 6:eabb5614.

42. Wang Z-H, An Y, Zhao T, Pei T-T, Wang DY, Liang X, Qin W, Dong T. 2025. Amidase and lysozyme dual functions in TseP reveal a new family of chimeric effectors in the type VI secretion system. eLife 13:RP101125.

43. Azhieh A, Hernandez P, Anderson AC, Sychantha D, Verster AJ, Whitney JC, Ross BD. 2025. Rapidly evolving orphan immunity genes protect human gut bacteria from intoxication by the type VI secretion system. bioRxiv 10.1101/2025.05.03.651265.

44. Kostiuk B, Santoriello FJ, Diaz-Satizabal L, Bisaro F, Lee K-J, Dhody AN, Provenzano D, Unterweger D, Pukatzki S. 2021. Type VI secretion system mutations reduced competitive fitness of classical Vibrio cholerae biotype. Nat Commun 12:6457.

45. Caplan J, Padmanabhan M, Dinesh-Kumar SP. 2008. Plant NB-LRR Immune Receptors: From Recognition to Transcriptional Reprogramming. Cell Host & Microbe 3:126–135.

46. Ng A, Xavier RJ. 2011. Leucine-rich repeat (LRR) proteins: Integrators of pattern recognition and signaling in immunity. Autophagy 7:1082–1084.

47. Yadav SK, Magotra A, Ghosh S, Krishnan A, Pradhan A, Kumar R, Das J, Sharma M, Jha G. 2021. Immunity proteins of dual nuclease T6SS effectors function as transcriptional repressors. EMBO Rep 22:EMBR202051857.

48. Abramson J, Adler J, Dunger J, Evans R, Green T, Pritzel A, Ronneberger O, Willmore L, Ballard AJ, Bambrick J, Bodenstein SW, Evans DA, Hung C-C, O’Neill M, Reiman D, Tunyasuvunakool K, Wu Z, Žemgulytė A, Arvaniti E, Beattie C, Bertolli O, Bridgland A, Cherepanov A, Congreve M, Cowen-Rivers AI, Cowie A, Figurnov M, Fuchs FB, Gladman H, Jain R, Khan YA, Low CMR, Perlin K, Potapenko A, Savy P, Singh S, Stecula A, Thillaisundaram A, Tong C, Yakneen S, Zhong ED, Zielinski M, Žídek A, Bapst V, Kohli P, Jaderberg M, Hassabis D, Jumper JM. 2024. Accurate structure prediction of biomolecular interactions with AlphaFold 3. Nature 630:493–500.

49. Osdaghi E, Abachi H, Jacques M-A. 2025. Clavibacter michiganensis Reframed: The Story of How the Genomics Era Made a New Face for an Old Enemy. Molecular Plant Pathology 26:e70093.

50. Osdaghi E, Taghavi SM, Hamidizade M, Kariminejhad M, Fazliarab A, Hajian Maleki H, Baeyen S, Taghouti G, Jacques M-A, Van Vaerenbergh J, Portier P. 2024. Multiphasic investigations imply transfer of orange-/red-pigmented strains of the bean pathogen Curtobacterium flaccumfaciens pv. flaccumfaciens to a new species as C. aurantiacum sp. nov., elevation of the poinsettia pathogen C. flaccumfaciens pv. poinsettiae to the species level as C. Poinsettiae sp. nov., and synonymy of C. albidum with C. citreum. Systematic and Applied Microbiology 47:126489.

51. Gonçalves RM, Balbi-Peña MI, Soman JM, Maringoni AC, Taghouti G, Fischer-Le Saux M, Portier P. 2019. Genetic diversity of Curtobacterium flaccumfaciens revealed by multilocus sequence analysis. Eur J Plant Pathol 154:189–202.

52. Kirchberger PC, Unterweger D, Provenzano D, Pukatzki S, Boucher Y. 2017. Sequential displacement of Type VI Secretion System effector genes leads to evolution of diverse immunity gene arrays in Vibrio cholerae. Sci Rep 7:45133.

53. Jurėnas D, Grandjean MM, Rudgalvyte M, Delobelle T, Elomari M, Marchal N, Oms T, Paillat M, Dieu M, Cascales E, Varoquaux N, Abby S, Fronzes R, Doan T. 2026. Rhs toxin complexes secreted via the type IX secretion system mediate kin discrimination in environmental Bacteroidota. Research Square 10.21203/rs.3.rs-9651787/v1.

54. Vassallo CN, Troselj V, Weltzer ML, Wall D. 2020. Rapid diversification of wild social groups driven by toxin-immunity loci on mobile genetic elements. ISME J 14:2474–2487.

55. Kobayashi K. 2021. Diverse LXG toxin and antitoxin systems specifically mediate intraspecies competition in Bacillus subtilis biofilms. PLOS Genetics 17:e1009682.

56. Liu M, Wang H, Wang Z, Wang H, Zhang K, Xue J, Liu R, Liu Y, Xia P, Wang H, Kan B, Li Y, Li S, Fu Y. 2025. A Vibrio-specific T6SS effector reshapes microbial competition by disrupting Vibrio bioenergetics. Cell Host & Microbe 33:1146–1160.e8.

57. Egamberdieva D, Kucharova Z, Davranov K, Berg G, Makarova N, Azarova T, Chebotar V, Tikhonovich I, Kamilova F, Validov SZ, Lugtenberg B. 2011. Bacteria able to control foot and root rot and to promote growth of cucumber in salinated soils. Biol Fertil Soils 47:197–205.

58. Egamberdieva D, Jabborova D, Berg G. 2016. Synergistic interactions between Bradyrhizobium japonicum and the endophyte Stenotrophomonas rhizophila and their effects on growth, and nodulation of soybean under salt stress. Plant Soil 405:35–45.

59. Alavi P, Starcher M, Zachow C, Mueller H, Berg G. 2013. Root-microbe systems: the effect and mode of interaction of Stress Protecting Agent (SPA) Stenotrophomonas rhizophila DSM14405T. Front Plant Sci 4.

60. Liu Y, Gao J, Wang N, Li X, Fang N, Zhuang X. 2022. Diffusible signal factor enhances the saline-alkaline resistance and rhizosphere colonization of *Stenotrophomonas rhizophila* by coordinating optimal metabolism. Science of The Total Environment 834:155403.

61. Sawa J, Malet H, Krojer T, Canellas F, Ehrmann M, Clausen T. 2011. Molecular Adaptation of the DegQ Protease to Exert Protein Quality Control in the Bacterial Cell Envelope *. Journal of Biological Chemistry 286:30680–30690.

62. Sprouffske K, Wagner A. 2016. Growthcurver: an R package for obtaining interpretable metrics from microbial growth curves. BMC Bioinformatics 17:172.

63. Blum M, Andreeva A, Florentino LC, Chuguransky SR, Grego T, Hobbs E, Pinto BL, Orr A, Paysan-Lafosse T, Ponamareva I, Salazar GA, Bordin N, Bork P, Bridge A, Colwell L, Gough J, Haft DH, Letunic I, Llinares-López F, Marchler-Bauer A, Meng-Papaxanthos L, Mi H, Natale DA, Orengo CA, Pandurangan AP, Piovesan D, Rivoire C, Sigrist CJA, Thanki N, Thibaud-Nissen F, Thomas PD, Tosatto SCE, Wu CH, Bateman A. 2025. InterPro: the protein sequence classification resource in 2025. Nucleic Acids Res 53:D444–D456.

64. Holm L. 2020. DALI and the persistence of protein shape. Protein Science 29:128–140.

65. Schindelin J, Arganda-Carreras I, Frise E, Kaynig V, Longair M, Pietzsch T, Preibisch S, Rueden C, Saalfeld S, Schmid B, Tinevez J-Y, White DJ, Hartenstein V, Eliceiri K, Tomancak P, Cardona A. 2012. Fiji: an open-source platform for biological-image analysis. Nat Methods 9:676–682.

